# AFMfit : Deciphering conformational dynamics in AFM data using fast nonlinear NMA and FFT-based search

**DOI:** 10.1101/2024.06.03.597083

**Authors:** Rémi Vuillemot, Jean-Luc Pellequer, Sergei Grudinin

## Abstract

Atomic Force Microscopy (AFM) offers a unique opportunity to study the conformational dynamics of proteins in near-physiological conditions at the single-molecule level. However, interpreting the two-dimensional molecular surfaces of multiple molecules measured in AFM experiments as three-dimensional conformational dynamics of a single molecule poses a significant challenge. Here, we present AFMfit, a flexible fitting procedure that deforms an input atomic model to match multiple AFM observations. The fitted models form a conformational ensemble that unambiguously describes the AFM experiment. Our method uses a new fast fitting algorithm based on the nonlinear Normal Mode Analysis (NMA) method NOLB to associate each molecule with its conformational state. AFMfit processes conformations of hundreds of AFM images of a single molecule in a few minutes on a single workstation, enabling analysis of larger datasets, including high-speed (HS)-AFM. We demonstrate the applications of our methods to synthetic and experimental AFM/HS-AFM data that include activated factor V and a membrane-embedded transient receptor potential channel TRPV3. AFMfit is an open-source Python package available at https://gricad-gitlab.univ-grenoble-alpes.fr/GruLab/AFMfit/.

## Introduction

Observing biomolecules at work in their native environment is crucial for understanding their biological function and developing therapeutic applications. Only a handful of experimental methods (such as SAXS or solution NMR) can provide this information at the atomic scale. Among these, Atomic Force Microscopy (AFM) is a unique imaging technique that captures snapshots of biomolecules at nanometer resolution in near-physiological conditions with a signal-to-noise ratio (SNR) high enough to see individual molecules. In the AFM experiments, one scans the molecules with a sharp tip that measures the topography of their surface. The surface profiles are not resolved sufficiently to observe atomic details, but they inform on the domain-level arrangement of single molecules in solution. Moreover, one can collect AFM recordings successively with a frame rate inferior to one second in high-speed (HS)-AFM. This technique has been widely used to observe various biomolecules at work [1–3].

One of the main challenges in AFM is the interpretation of the resolution-limited 2D topographic surfaces of flexible biomolecules into understandable conformational dynamics in 3D. Although the direct reconstruction of a 3D contour of the molecule and its dynamics from the 2D surfaces is theoretically possible in case of sufficient observations, it has severe limitations. Firstly, the molecule in an AFM experiment needs to be attached to a flat surface of the stage to be imaged and tends to adopt the same orientation. Such a preferred viewing direction would cause anisotropy in the 3D reconstruction, a well-known problem in cryo-EM [4]. Secondly, the spatial resolution in AFM imaging is typically limited by the size of the tip to the nanometer scale [5]. Therefore, the AFM images are resolved only to the domain level, and atomic-level dynamics are not directly observable and would require prior structural knowledge.

Due to these limitations, computational methods to interpret AFM data in 3D typically use a model-based approach [6–16]. In such an approach, one or several available 3D structures at atomic resolution, such as those available in the Protein Data Bank (PDB), are docked under the AFM topographic images to estimate their position, orientation, and, possibly, their conformation. Most available computational methods focus on the rigid-body fitting of one or a few static candidate structures to AFM images [8, 10–13]. These candidate structures are, typically, stable conformations solved by X-ray crystallography or cryo-EM. These methods reconstruct the conformational variability using multiple molecular states that must be preliminary available to the user. This requirement has several limitations. Firstly, there is no guarantee that a sufficient number of states is available to describe the data. Secondly, due to crystallographic contacts, the states solved in crystals may not correspond to those observed in solution when probed by the AFM. Finally, the flexibility of biomolecules is typically better described by continuous motions rather than a few discrete states. One can partly avoid these limitations by enriching the number of candidate structures by sampling new states with molecular dynamics (MD) simulations [11, 12] at the expense of sufficiently long simulation trajectories that cover all the possible states encountered in the data. Another way to infer the flexibility of biomolecules is by fitting multiple rigid structures corresponding to the macromolecular subunits under study to the AFM images, allowing rigid-body motions between the subunits [7, 9].

The problem of reconstructing the continuous flexibility of biomolecules, as opposed to discrete states, from heterogeneous 2D views has been widely studied in cryo-EM and has undergone extensive developments in recent years [17–31]. In particular, continuous-state approaches emerged, which intend to treat each single molecule (designated as a particle in the cryo-EM field) as a different conformation and try to place each particle onto a continuous conformational space. This space typically contains 10^5^ to 10^6^ data points corresponding to each particle and gives an overall understanding of the conformational distribution in the sample. Some recent methods use model-based approaches that infer molecular flexibility using computational simulations from an initial atomic model, including MD simulations [21, 22, 29], the normal mode analysis (NMA) [17, 27, 28], and Zernike polynomials [25].

One can apply a similar approach to AFM. Recently, researchers have proposed two methods to infer the conformational state from a single-molecule AFM image using molecular simulations. One method uses a coarse-grained model represented as a mixture of Gaussians and reconstructs the structural heterogeneity with Monte Carlo simulations constrained by AFM data [14, 16]. The other method performs flexible fitting of an initial model using MD simulations biased with AFM-derived forces [15]. These methods have been applied to describe the conformation of one or a few particles (up to twelve [16]) but have not been tested on larger datasets.

However, AFM images often contain hundreds to thousands of single molecules, and this number may increase even further when recording multiple frames, such as in HS-AFM. Processing a larger number of images can result in a more accurate and comprehensive description of the conformational dynamics in the sample, in particular in the context of continuous motions of highly flexible biomolecules.

In the present work, we propose AFMfit, a new method for reconstructing conformational dynamics from AFM experiments. Our algorithm uses a novel flexible fitting procedure that scales to many single molecules in AFM images. AFMfit explores the flexible degrees of freedom with the nonlinear NMA method NOLB [32]. Our approach has the advantage of being very fast (e.g., a few tens of minutes on a personal desktop), requiring only one atomic model, and avoiding the structural distortions typically induced by linear normal modes at large amplitudes [32]. The method outputs a conformational landscape that describes the principal structural variations and their distribution over the data. We present our approach and demonstrate its applications to synthetic and experimental AFM and HS-AFM data. We show that the method accurately predicts the conformational dynamics given a set of noisy synthetic AFM images. Then, we apply the method to experimental an AFM dataset of activated coagulation factor V (FVA) [33] and an HS-AFM dataset of a transient receptor potential, TRPV3 [34]. The results highlight the conformational dynamics of the studied biomolecules and corroborate the previously published findings [33, 34].

## Results

### AFMfit’s rigid and flexible fitting

AFMfit aims to find the conformational ensemble of the sample molecule that best describes a set of AFM topographic images. More specifically, for each image in the set, the algorithm searches for the best rigid (3D rotation and translation) and flexible (amplitudes of the normal mode deformations) alignments of a given initial atomic model that maximizes its similarity with the experimental data.

We conduct the rigid and flexible fitting procedures in the image space, meaning that we compute 2D AFM-like projections of the initial 3D model and compare them with the raw experimental observations to find the best match. In the first step of our procedure, we perform a global rigid fitting to estimate the global orientation of the molecule in each AFM image. Then, a flexible fitting algorithm based on nonlinear normal modes [32] locates the best flexible deformations of the initial model to describe the image, given the global rigid alignment found in the previous step. Note that the flexible fitting is also associated with local continuous rigid alignment, which allows for rigid rearrangement during the fitting. Finally, the deformations associated with each image form a conformational ensemble. We can further interpret this ensemble by projecting it onto a low-dimensional space of conformations using Principal Component Analysis (PCA). Figure 1 schematically shows the fitting procedure.

**Fig. 1.**
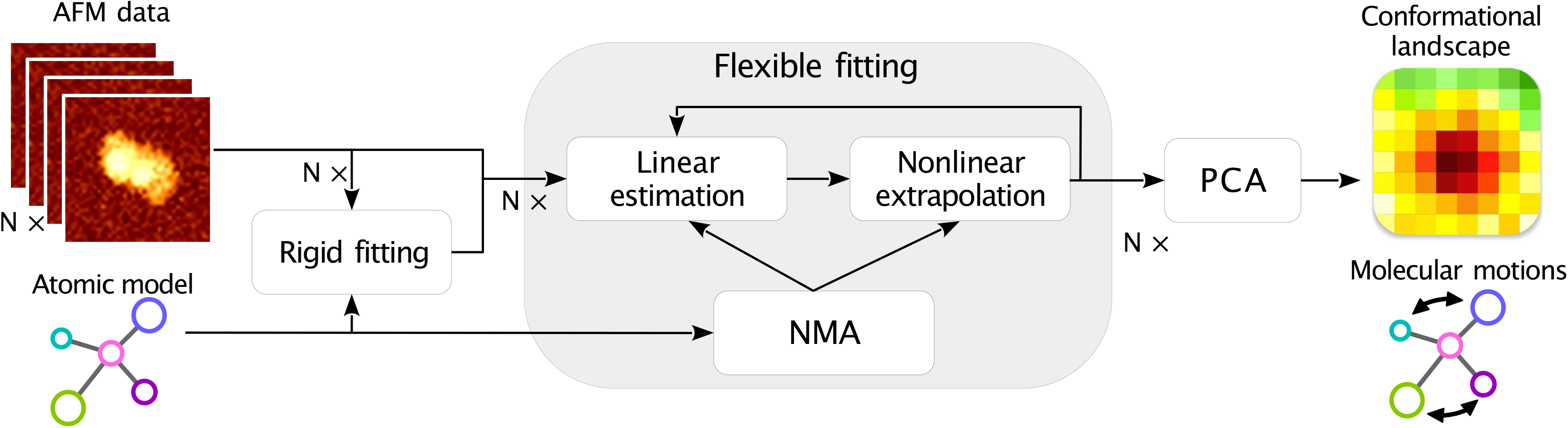
Schematic workflow of the AFMfit fitting procedure. The method takes as input an available atomic model and a set of *N* AFM images of single molecules. The fitting consists of three main steps. In the first step, we perform a global rigid fitting of the atomic model to each AFM image. In the second step, we determine a set of continuous flexible and rigid deformations that improve the fit. The flexible deformations are obtained by performing NMA, and the resulting normal modes are fit in an iterative two-step procedure. Firstly, the optimal set of deformations along the linear normal modes is estimated. Secondly, a non-linear extrapolation of the deformations is carried out following the NOLB method [32] that preserves the model structure. In the last step, we assemble the flexibly fitted models into a low-dimensional representation of the conformational ensemble and extract molecular motions.

### Evaluation of AFMfit on synthetic data of EF2

As a first step in evaluating our technique, we created a synthetic AFM data to test its performance in a controlled environment. We used a protein model of Elongation Factor 2 (EF2) that is available in the PDB (code 1N0V) [36]. This protein is suitable for testing our fitting pipeline because it has an asymmetric structure that consists of several globular domains, shown in Fig. 2a. When we deform these domains with NMA, they could adopt a cylinder-like shape with a pseudo-symmetry axis, which makes the fitting ambiguous at the AFM resolution. This ambiguity makes the rigid fitting challenging and requires the fitting pipeline to disentangle the conformation and the orientations to achieve a good fit. We created a conformational ensemble of 100 conformations by applying flexible deformations to the EF2 model using the nonlinear normal mode analysis method NOLB [32]. We used the first two nonzero-frequency normal modes (number 7 and 8 shown in Fig. 2a) to generate these deformations. We selected the protein conformations along these deformations to be on average of 4 Å RMSD and describe a circle in the conformational space obtained through PCA on the conformational ensemble (as shown in Fig. 2c). We then applied 3D rotations to the conformational ensemble so that the models lie approximately flat on the *xy* - plane. We then generated pseudo-AFM images from the conformational ensemble of 40 × 40 pixels with a pixel size of 7 Å using the collision-detection method of the “afmize” software [37]. We set the simulated probe angle to 10° and the probe radius to 2 nm. We added Gaussian noise with a standard deviation of 3 nm to match the typical SNR of experimental AFM data. It is important to note that this simulation method is conceptually different from the one used in the flexible fitting pipeline to recover the flexible deformations. We used AFMfit with a 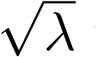 value of 5, a *σ* value of 4.2 and included the five lowest-frequency normal modes in the fitting.

**Fig. 2.**
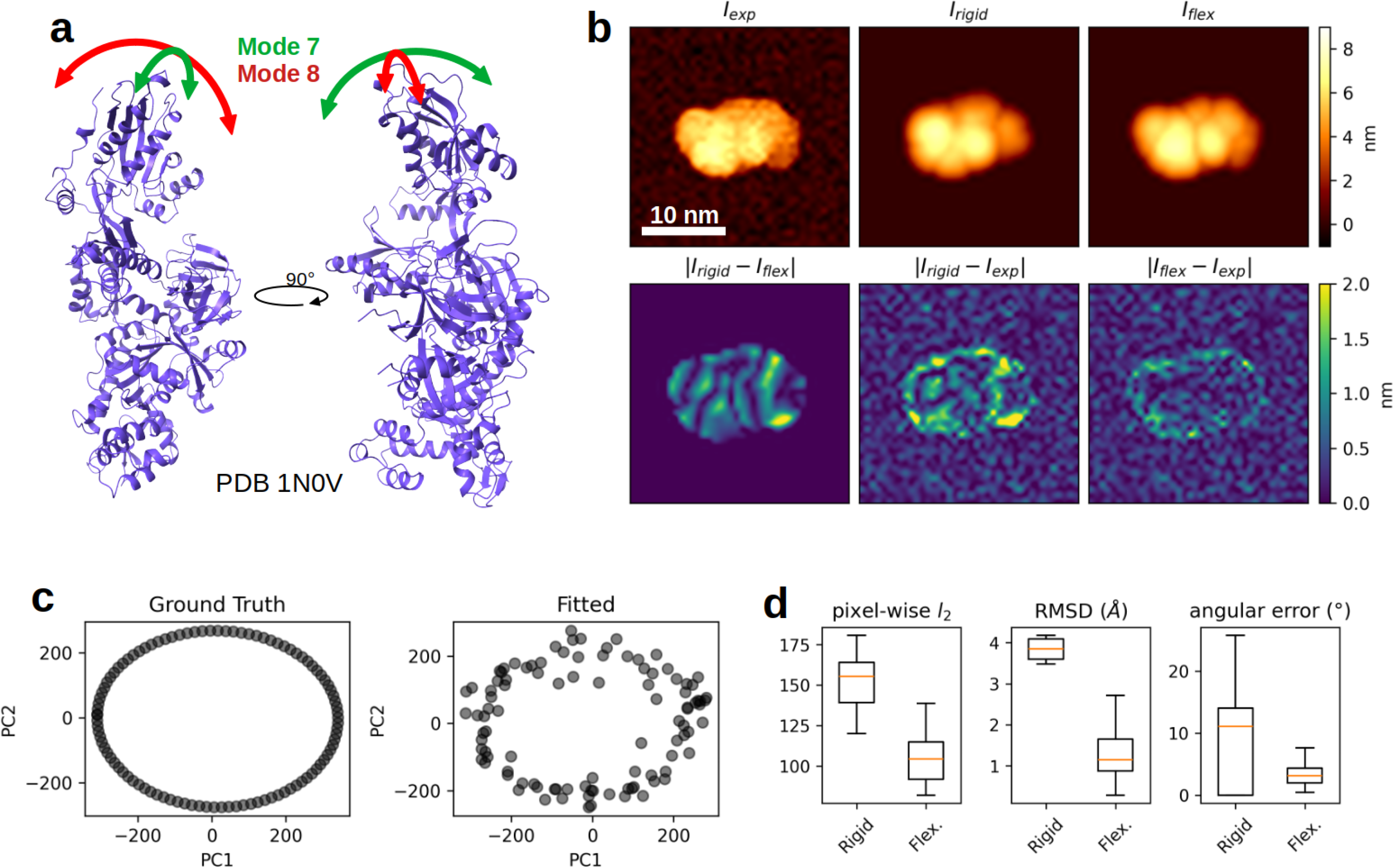
AFMfit applied to synthetic data of EF2. **a**. 3D atomic structure of EF2 (PDB 1N0V), the arrows show the motion along the normal modes used to generate the synthetic dataset (modes 7 and 8). Molecular visualizations: ChimeraX [35]. **b**. Fitting results for a selected particle designated as *I*_*exp*_. The best-fitted projections of the rigid and the flexible fitting are designated as *I*_*rigid*_, and *I*_*flex*_, respectively. The second row shows the absolute value of the difference (in nm) between *I*_*rigid*_, *I*_*flex*_ and *I*_*exp*_. **c**. Conformational space obtained by PCA on the ground truth (left) and the fitted (right) atomic models. **d**. Statistics of the flexible fitting compared to the rigid fitting, from left to right, the pixel-wise *l*_2_ norm used to drive the fitting, the RMSD between the ground truth and the fitted atomic models, and the angular error between the ground truth and the estimated particle orientations.

AFMfit was able to recover the ground truth conformational ensemble from the synthetic AFM data of EF2. Figure 2b shows an example of fitting for one selected particle in the set. The top left panel shows the particle, and the top middle and top right panels demonstrate the best match for rigid and flexible fits, respectively. If we examine this selected example, we observe that the rigid fitting alone misalignes the model with the image. Indeed, the best orientation obtained by the rigid fitting has an angular error close to 180° compared to the ground truth, meaning that the model flipped around the pseudo-symmetry axis. However, the flexible fitting corrected the misalignment and found the correct conformation, reducing the angular error to 3° (see the differences between rigid and flexible fitting, bottom row of Fig. 2b).

We further analyzed the fitted conformational ensemble (together with the ground-truth ensemble) with PCA. Figure 2c shows its first two components, with the left panel showing the samples from the ground truth and the right panel – the ones from the fitted ensemble. We can observe that both the fitted and the ground-truth ensembles closely match in terms of the amplitude of the deformations and the distribution of the conformations, as seen by the circle shape in the first two principal components.

Figure 2d provides a further comparison between the rigid and flexible fits. We found that the pixel-wise *l*_2_, used to drive the fitting, was significantly decreased between the rigid and flexible fitting procedures, with an average decrease of 34% from its initial value. Additionally, the RMSD between ground truth and fitted structures decreases from an average of 3.9 Å to 1.1 Å, and the angular error between the ground-truth orientation and the fitted orientation decreases from an average of 11.1° to below 3.1°. Overall, we can conclude that the presented method successfully recovered the correct conformational ensemble from a set of noisy synthetic AFM images of EF2.

### Analysis of experimental AFM data of FVa

In the second stage of our method assessment, we applied the fitting pipeline to experimental AFM data of activated factor V (FVa). FVa is a protein involved in the blood coagulation cascade. In 2014, Chaves et al. obtained a high-resolution AFM measurement of isolated human FVa in a liquid environment containing multiple particles of FVa [33].

FVa is a heterodimer composed of a heavy chain with two A domains (A1-A2) and a light chain with one A domain and two C domains (A3-C1-C2). The C domains are known to be implicated in the binding of FVa with phospholipid membranes, which is crucial to its function, as shown by mutagenesis studies [38, 39]. The 2014 study revealed dynamic variations of the C1 and C2 domains [33]. At the time of the study, the structure of full human FVa was unknown. To assemble crystal structures of A trimers and two of C domains, the authors used a computational reconstruction pipeline known as AFM-assembly [7]. They reported conformational changes in the angle formed by C1 and C2 domains on a selected set of 20 particles [33].

More recently, Ruben et al. derived a complete structural model of human FVa (PDB code 7KXY) from a cryo-EM map of the Prothrombin-FVa-FXa complex at 4.1 Å resolution [40]. We used this model as the initial state to apply our fitting algorithm to the AFM data of FVa. We performed NMA and selected the six zero-frequency and eight lowest-frequency (nonzero) normal modes, which mainly correspond to motions of the C domains. We increased the number of extracted particles in the dataset from 20 to 77 by a manual selection and applied a masking operation composed of low-pass filtering and threshold to remove most of the background noise.

### AFMfit recovers dynamics of the C domains of FVa

Overall, the fitting procedure decreased the loss function significantly by an average of 7.4 % between the rigid and the flexible fitting. Figure 3a-b shows a comparison between the rigid and flexible fitting results for a selected particle from the dataset. In this example, we observe that the C2 domain of FVa rotates (see red arrows in Figure 3a-b), decreasing the pixel-wise *l*_2_ by 18.5 % (see bottom panels of Figure 3a).

**Fig. 3.**
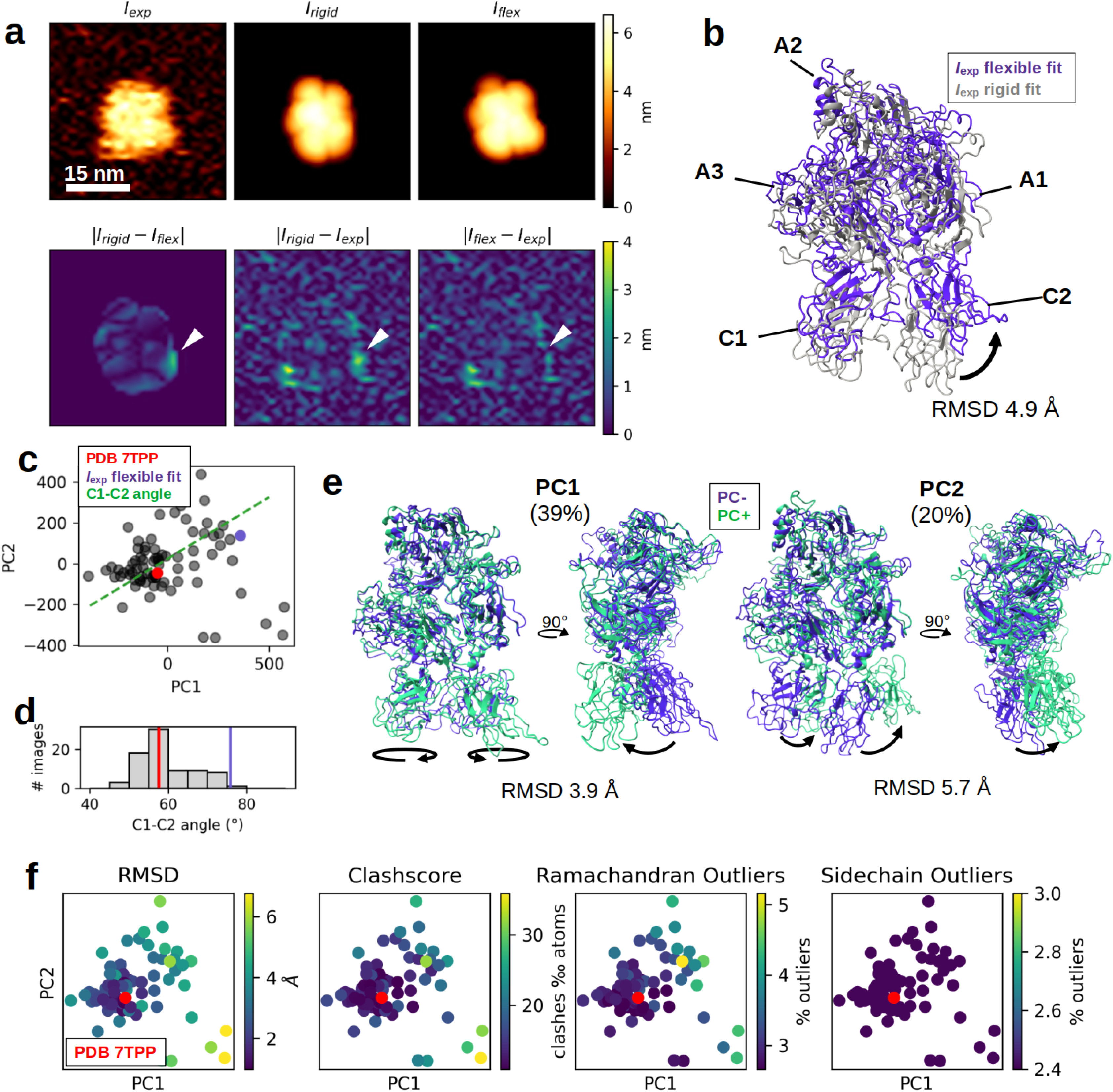
AFMfit applied to the experimental AFM data of FVa. **a**. Fitting results for a selected AFM image designated as *I*_*exp*_. The best-fitted projection of the rigid and the flexible fitting are designated as *I*_*rigid*_ and *I*_*flex*_, respectively. The second row shows the absolute value of differences (in nm) between *I*_*rigid*_, *I*_*flex*_, and *I*_*exp*_. The arrow heads show the location of C2. **b**. 3D structure of the best-fitted atomic models for *I*_*exp*_ (see panel (a.)) for the rigid fitting, PDB 7TPP (grey) and the flexible fitting (blue). **c**. Conformational space obtained by PCA on the set of flexibly-fitted atomic models. The first two principal components are shown, the red dot corresponding to the location of the initial model and the purple dot corresponding to the flexibly fitted model for the selected image *I*_*exp*_. The green dashed line corresponds to the C1-C2 angle motion direction obtained by linear regression of the C1-C2 angle with respect to PC1 and PC2. **d**. Histogram of the C1-C2 angles of the flexibly-fitted atomic models. The red line corresponding to the C1-C2 angle of the initial model and the purple line corresponding to the fitted model for *I*_*exp*_. **e**. The conformational changes associated with PC1 (39 % of explained variance) and PC2 (20 % of explained variance) are shown as a superposed atomic model associated with positive (purple) and negative (green) motions along the respective principal components. The arrows outline the conformational changes. **f**. Structural quality measures are shown on the first two principal components. From left to right, the RMSD between the initial and the fitted models, the clashscore corresponding to the number of steric clashes per 1000 atoms, the Ramachandran outliers corresponding to non-favorable dihedral angles, and the sidechain outliers corresponding to non-favorable rotamers. The red dots correspond to the location of the initial model.

We further processed the 77 fitted models with PCA to obtain the conformational space shown in Figure 3c. The first two principal components account for 60 % of the total variance (39 % for PC1 and 20 % for PC2). The conformational changes associated with PC1 and PC2 involve motions of the C domains, as shown in Fig.3e. For PC1, the C domains are displaced collinearly to the pseudo-symmetry axis formed by the A trimer (Supplementary Video 1). The conformational change described by PC1 is also associated to a rotation of C1 and C2 that opens the angle between the two C domains. For PC2, the C domains rotate in the plane formed by the A trimer 0(Supplementary Video 2).

We note that the motion associated with a the combination of PC1 and PC2 reassembles the reconstructed models from the assembly protocol in [33]. More precisely, in the 2014 study, the authors reported a variation in the angle formed by the C1 and C2 domains. Here, the two most dominant reconstructed motion are associated with a variation of the angle between C1 and C2. Figure 3d shows the distribution of the C1-C2 angle over the fitted models. The distribution closely matches a linear combination of PC1 and PC2 (green dashed line in Figure 3c).

### AFMfit conserves the structural quality of the initial model

To ensure that the produced conformational ensemble conserves the structural elements of the initial model, we calculated several standard metrics for the structural quality over the entire ensemble. We selected metrics similar to those used to assess cryo-EM-derived models in the PDB: the *clashscore* that measures the number of steric clashes (including hydrogens) per 1000 atoms, the *Ramachandran outliers* that count the percentage of disallowed dihedral angles and the *sidechain outliers*, and the percentage of disallowed rotamers. We computed all the metrics using the MolProbity software [41].

We shall note that the initial model of FVa used in this study was reconstructed from a medium-resolution (4.1 Å) EM map with some regions poorly resolved. Therefore, some of these metrics are initially of average quality, and the objective of using these metrics is not to give an absolute value of the quality of the model but rather to ensure that the metrics remain consistent and the quality does not deteriorate.

Figure 3d shows the value of these metrics, together with the RMSD between the initial and fitted models (first panel) projected on the two first principal components. For low to medium RMSD values (below 6 Å), corresponding to 93 % of the ensemble, the three metrics are close to the initial values, suggesting that the structural quality of the initial model remains stable. For high RMSD values (above 6 Å), which account for solely 7 % of the ensemble, the structural quality decreases (clashscore above 30 ‰, Ramachandran outliers above 5 %). We shall expect this behavior, as using normal modes to approximate molecular deformations is valid only in the close vicinity of the equilibrium conformation (i.e., the initial model, in this case).

### AFMfit recovers compositional and structural heterogeneity from HS-AFM movies of TRPV3

In the last stage of assessment of the method, we processed HS-AFM movies of a membrane-embedded transient receptor potential channel, TRPV3 [34]. We processed two movies of TRPV3 present in two oligomeric states, pentameric and tetrameric. The first HS-AFM movie contains multiple TRPV3 single molecules that show dynamic transitions between two oligomeric states. The second movie has a higher magnification and captures two single molecules at high resolution, one in the pentameric state and the other in the tetrameric state.

Apart from the apparent flexibility, the movies contain compositional heterogeneity (different oligomeric states). As AFMfit relies on the flexible deformation of an initial model, by default, it does not consider compositional heterogeneity. To tackle this issue, we used two initial models (TRPV3 pentamer, PDB code 8GKG; TRPV3 tetramer, PDB code 8GKA) and carried out the AFMfit protocol over all the particles with the two models. To assign the best model to each single molecule, we selected those that achieved the highest correlation on each particle.

We analyzed the first HS-AFM movie and identified up to 13 single molecules over 27 frames, with a maximum of 10 single molecules in one frame present simultaneously, resulting in 172 single-molecule frames. The AFMfit results, starting from the two oligomeric states (PDB codes 8GKG and 8GKA), allowed us to identify that 11 % of the molecules were pentamers. Supplementary Video 3 shows the result of the fitting for the entire movie and Figure 4a shows the result for one selected frame.

**Fig. 4.**
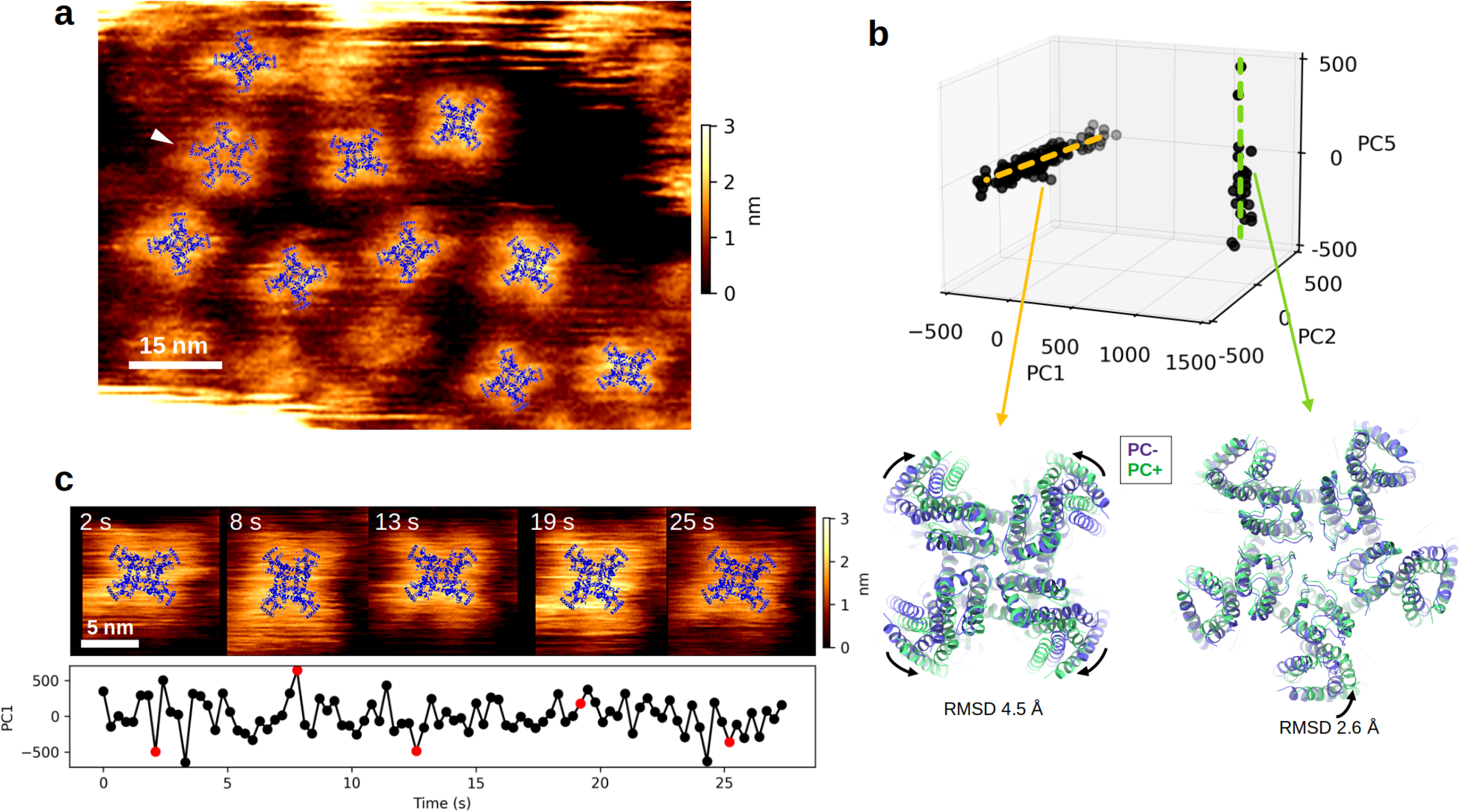
AFMfit applied to HS-AFM data of TRPV3. **a**. An example of a fitted HS-AFM frame with several pentameric states of TRPV3 (arrowheads). The models fitted by AFMfit are superimposed with the HS-AFM frame for each single molecule. **b**. The first three components of the PCA decom-position from the 255 models fitted by AFMfit on the HS-AFM movie are shown in (a.). The first components separate pentameric and tetrameric states. The second component is mainly associated with motions in the pentameric state (green dashed line); the corresponding structural changes are shown in the right-most structure. The third component is mainly associated with motions in the tetrameric state (yellow dashed line); the corresponding structural changes are shown in the left-most structure. **c**. Flexible motions observed in high-magnification HS-AFM movie of TRPV3 in the tetrameric state. The top row shows the relevant HS-AFM frame superimposed with the corresponding models fitted with AFMfit. The bottom row shows the time series of the state along the first principal component (red dots correspond to the frames shown in the top row).

Apart from the compositional heterogeneity, we analyzed the flexibility of each single molecule in the first HS-AFM movie with PCA over the fitted models. Figure 4b shows the first three principal components of the 255 fitted models. We can see that the first component (80 % of explained variance) separates the two oligomeric states. The second, third, and fourth components (8.1 %, 4.4 % and 2.5 % of explained variance, respectively) are associated with flexible motions of the tetramers. The fifth component is associated with flexible motions of the pentamer (1.5 % of explained variance). The most dominant flexible motion for the tetramer (second principal component) is an inplane flattening (Figure 4b left and Supplementary Video 4) and for the pentamer (fifth principal component) is an asymmetric up and down motion in the *z*-axis (Figure 4b right and Supplementary Video 5). We should note that these flexible motions are relatively low-amplitude, especially for the pentamer (RMSD 4.3 Åand 2.9 Åbetween extremums, respectively).

We analyzed these motions further in the second HS-AFM movie at a higher magnification (pixel size of 1.25 Å). The movie contains 92 frames with a sampling rate of 0.3 seconds with two single molecules with stable oligomeric states (one pentamer and one tetramer). Supplementary Video 6 shows the result of the fitting for this movie.

The flexible motions extracted from these two single molecules are the same as the flexible motions described for the first movie (Figure 4b), i.e., small-amplitude inplane flattening for the tetramer and up and down motions of the monomers in the z-axis for the pentamer. Figure 4c shows the fitting results for the tetramer TRPV3 of the second movie over time. We selected relevant states at different locations along the main flexible component (PC1). A positive displacement along PC1 corresponds to an inplane flattening of the TRPV3 tetramer in the x-axis direction, and a negative displacement corresponds to the same motion in the y-axis direction. Most interestingly, there is no apparent time coordination between these small-amplitude flexible motions. Overall, we conclude that the AFMfit protocol separated the different oligomeric states in the HS-AFM movies of TRPV3 and assigned small-amplitude flexible deformations to each state.

## Discussion

One advantage of AFMfit is its speed, which allows the processing of hundreds to thousands of AFM images in just a few minutes on a personal desktop. Table 1 reports the computational time to run the AFMfit experiments presented in this work.

**Table 1.** Execution time of AFMfit for the presented AFM/HS-AFM datasets on a single workstation Intel(R) Core(TM) i7-13700 CPU @ 5.2 GHz.

| Dataset | # image | Pixel size (Å) | Time (min) |
| --- | --- | --- | --- |
| EF2 | 100 | 7.0 | 15 |
| FVa | 77 | 9.75 | 19 |
| TRPV3 (1<sup>st</sup>) | 172 | 4.0 | 9 |
| TRPV3 (2<sup>nd</sup>) | 184 | 1.25 | 23 |

To approximate low-dimensional motion manifolds of the input models, we employed NMA. We must notice, however, that using NMA for simulating the flexibility of biomolecules may have limitations. For example, one shortcoming of linear normal modes is that at high deformation amplitudes, the interatomic distances are no longer respected, and some parts of the molecular structure tend to get distorted. In our model, we have utilized a nonlinear NMA approach, NOLB, that brings a solution to this issue [32]. Another limitation of NMA is that its energy model predetermines the motions, and it can be challenging to pick the relevant ones. The elastic network model inside NOLB and many other NMA approaches accelerates the computations but may restrict the accuracy of the normal modes to the lowest frequencies. In our case, selecting a few low-frequency normal modes is suitable as the resolution of the AFM images is limited, and using more degrees of freedom would be prone to overfitting. Therefore, we recommend using less than ten lowest-frequency normal modes, and selecting the lowest ones would give acceptable results for most models. We shall note that we did not employ any restriction on the elastic network model regarding the interaction between the molecule and the surface to which it is attached. Such interactions can be included in a future analysis, as they may influence the flexibility of the structural model.

## Conclusion

To conclude, we propose a new fitting method to interpret conformational dynamics from AFM/HS-AFM using an input atomic model. To the best of our knowledge, it is the first approach that applies NMA-based fitting to the analysis of AFM images. We confirmed its ability to recover a given conformational space from synthetic data. We also tested the method on experimental datasets of AFM and HS-AFM and demonstrated that our results corroborate with previously published results [33, 34].

## Methods

### Simulating pseudo-AFM images

AFMfit compares experimental AFM images with pseudo-AFM images synthetically generated from the atomic models in different conformations and orientations. To simulate a pseudo-AFM image from a 3D atomistic model, a conventional approach consists of calculating the height in the *z* -direction at which a tip perpendicular to the *xy* -plane collides with the molecular structure, the so-called collision-detection method [10]. However, this method is not differentiable, which is a limitation for our flexible fitting purposes. Instead, we use a smooth differentiable function proposed in [15] that demonstrated a fair approximation to the collision-detection method.

The smooth function combines a set of Gaussian functions in the *xy*-plane that define the distance between each atom and the pixel center and a soft-maximum log-sum-exp function in the *z*-direction. Given an atomic model *X* composed of *N* atoms *X*_*i*_ = {*x*_*i*_, *y*_*i*_, *z*_*i*_}, *i* = 1, *N* , the smooth function П : 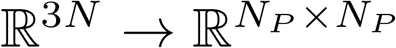 for the *p*th pixel *p* = {*p*_*x*_, *p*_*y*_} is given by :

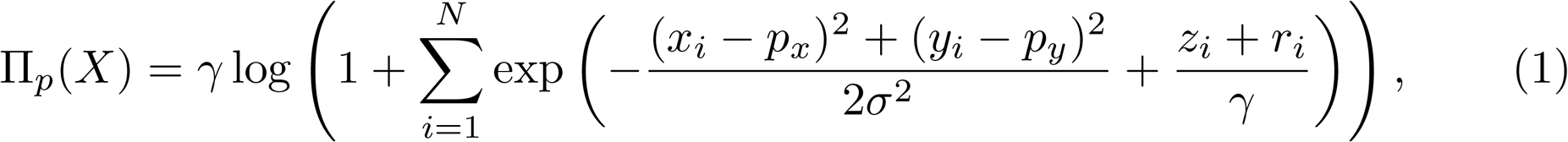

where *r*_*i*_ is the radius of the *i*th atom, *γ* is a parameter that makes the function Π_*p*_(*X*) smooth and *σ* is the spatial range (resolution) of the function. In our experiments, we use a fixed value of the resolution parameter *γ* = 1 Å. For this value of *γ*, the authors in [15] determined the optimal value of *σ* that best approximates the collision-detection method. This value depends on two parameters, the radius and the angle of the tip, and Table 1 in [15] describes the optimal *σ* given a range of values of these two parameters. We shall note that it can be challenging to accurately determine these parameters in the experimental setup. Therefore, this table provides a first estimate of *σ*, which can be adjusted later by running multiple experiments around this value and choosing the best fit. For the experiments, we used a value of *σ* of 3.2, 5.0 and 4.0 for the dataset of EF2, FVa and TRPV3, respectively.

### Global rigid fitting

The objective of the rigid-body fitting in AFMfit is to find the optimal rigid alignment (3D rotation and translation) of an atomic model *X* and the experimental AFM images preliminary to the flexible fitting. We will then refine the rotation and the translation obtained at this stage by the flexible fitting described below.

The algorithm exhaustively searches for the best rigid-body transformed model 𝒳 (*θ, τ* ) rotated by the 3D rotation described by the three Euler angles *θ* ∈ ℝ^3^ and translated by three translation parameters *τ* ∈ ℝ^3^ by minimizing the squared pixel-wise *l*_2_ norm between the experimental AFM image *Y* and a pseudo-AFM image generated from 𝒳 (*θ, τ* ) with the function described in Eq. 1 :

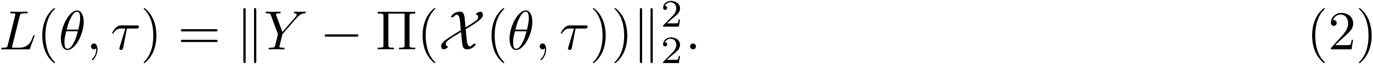

We use the pixel-wise *l*_2_ norm as the similarity measure (similarly as for the flexible fitting in Eq. 12). We have chosen the pixel-wise *l*_2_ over other measures (e.g., correlation coefficient widely used in cryo-EM), notably because it changes by uniform scaling Π(*X*) → *s*Π(*X*) and uniform shifting Π(*X*) → Π(*X*) + *s*. In comparison, the correlation coefficient is invariant to uniform scaling and uniform shifting. The changes to uniform shifting may help place the atomic model at the correct height in the topographic AFM image, and the changes to uniform scaling ensure the conservation of the geometric distances between molecular structures at different heights.

The algorithm exhaustively generates a pseudo-AFM image for discretized values of the three rotation parameters *θ* and one translation parameter *τ*_*z*_ (the translation in the *z* -direction). We accelerate the calculation of the last two translation parameters, translations in the *xy* -plane, via the fast Fourier transform calculation of *L* described later in this section.

To evenly discretize the space of 3D rotations given an angular increment Δ*θ*, we start by evenly distributing the viewing directions along the 3D sphere with a spherical Fibonacci lattice, which samples the two first Euler angles. Then, the remaining Euler angle describing an in-plane rotation (rotation about the *z* -axis) is discretized linearly with a Δ*θ* step. We linearly discretize the translation along the *z*-axis with an increment Δ*τ*_*z*_.

Finally, we obtain the last two translation parameters *τ*_*x*_ and *τ*_*y*_ (translation in the *xy* -plane) by an FFT-accelerated calculation of *L*. Concretely, we can rewrite *L* as

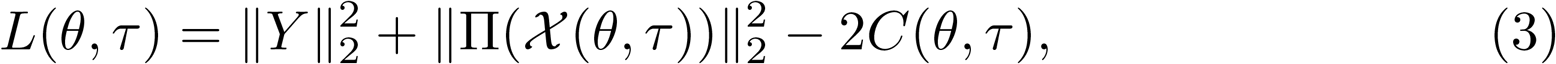

where

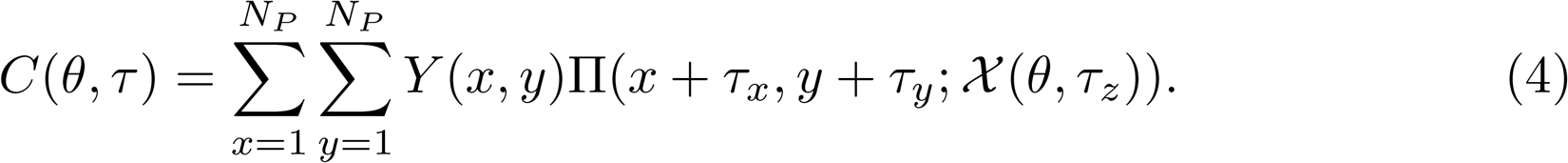

Note that we apply the translation in the *xy* -plane in Eq. 4 in the image space rather than the 3D Cartesian space, therefore *C*(*θ, τ* ) is also the 2D cross-correlation between *Y* and Π(*𝒳* (*θ, τ*_*z*_)). For a given value of *θ* and *τ*_*z*_, considering that 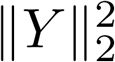 is constant and 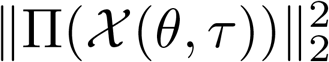 is invariant to translation in the *xy* -plane, the values of *τ*_*x*_, *τ*_*y*_ that maximize *C* also minimize *L* and can be calculated for all values of *τ*_*x*_ and *τ*_*y*_ in the discrete image space {1, … , *N*_*P*_ } via the fast Fourier transform :

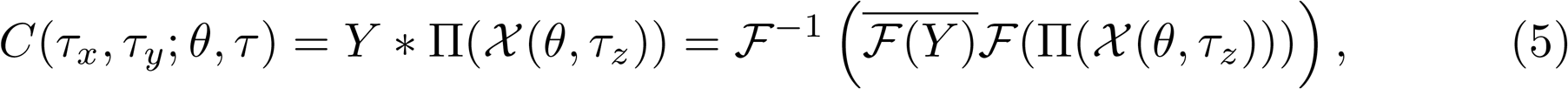

where the * sign denotes the 2D cross-correlation, ℱ is the 2D Fourier transform, ℱ^*−*1^ is the inverse 2D Fourier transform, and 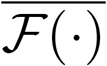 is its complex conjugate.

### Normal Mode Analysis and NOLB

The core behind the flexible fitting part of our method is a low-rank representation of molecular flexibility computed by the NOLB method [32]. In a nutshell, NOLB represents the molecular system using an all-atom elastic network model with an anisotropic potential function of the following form,

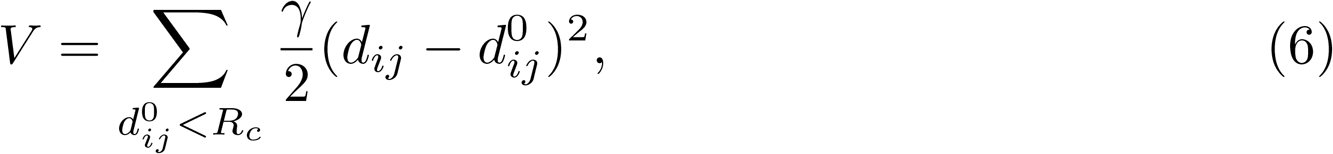

where *d*_*ij*_ is the distance between the *i*^*th*^ and the *j*^*th*^ atoms, 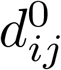 is the reference distance between these atoms, as found in the original model, *γ* is the stiffness constant, and *R*_*c*_ is a cutoff distance, which we fix to 8 Å by default. Then, using this potential function, it constructs a mass-weighted Hessian matrix and projects it into the space where each residue is represented as a rigid body. After, the method computes several lowest-frequency eigenvectors in the reduced rigid-residue space. We fix this number to a maximum of ten, and one can select the most relevant vectors manually, e.g., by visualizing each of them to get a finer description of the flexible motions. Note that using a higher number of normal modes is prone to overfitting. Each eigenvector has six components per residue: three correspond to the instantaneous linear velocities 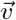 and the other three to the instantaneous angular velocities 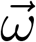. At the same time, we also compute an all-atom linearization *A* of the normal modes projecting the eigenvector back to the all-atom space. Finally, following the NOLB methodology, we extrapolate coordinates *X*(*α*) of each atom in a residue at an amplitude *α* as

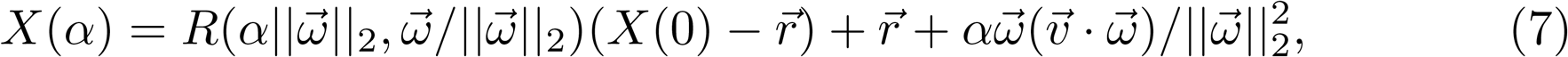

where we parameterize a rigid rotation *R* using an angle-axis notation. In this equation, the center of rotation 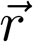 for a residue with the center of mass 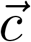 is given by

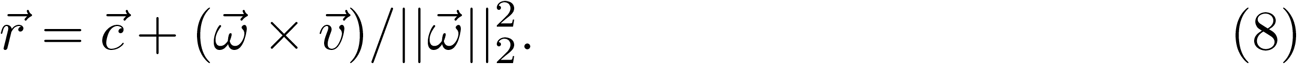

### Flexible fitting of an AFM image

Our flexible fitting procedure operates in two steps that are iterated until the algorithm convergence. In the first step, we estimate the optimal change in the linear normal mode amplitudes Δ*α* ∈ ℝ^*M*^ using *M* lowest frequency normal modes, such that

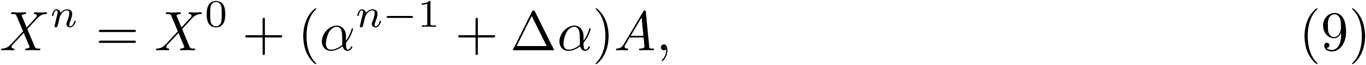

where *X*^*n*^ is the atomic model estimated at the *n*-th iteration of the algorithm, *α*^*n−*1^ ∈ ℝ^*M*^ are the normal mode amplitudes at the previous iteration, *X*^0^ is the known initial atomic model, and *A* ∈ ℝ^*M ×*3*N*^ is the set of *M* low-frequency normal modes associated to *X*^0^. In the second step, we update the normal mode amplitudes *α*^*n*^ = Δ*α* + *α*^*n−*1^ and extrapolate the atomic model 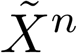 with the nonlinear NOLB procedure explained above. We then use the obtained model 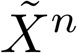 to calculate the change in normal mode amplitudes in the first step of the next iteration. Then, we repeat the two steps until convergence. Typically, the algorithm converges in less than ten iterations. Note that during the process, the normal modes *A* do not need to be updated and can remain constant, which reduces the computational cost.

We shall note that the set of *M* low-frequency normal modes includes the six modes *A*_*i*_ with *i* = {1, … , 6} with *zero* frequency corresponding to rigid-body translations and rotations. We include these six additional degrees of freedom in the fitting procedure to allow for continuous rigid-body rearrangement during the flexible deformations. These rigid rearrangements are crucial for the accuracy of our method, as the global rigid fitting might not be valid when the conformation of the model is changed.

We compute the *zero* frequency modes analytically according to [42]. The three modes associated with a rigid translation are straightforward to express as a displacement in the Cartesian space. The three other modes correspond to three infinitesimal rotations about the *x*,*y*, and *z* axes. We can compute an infinitesimal rotation *A*_*R*_(*u, θ*) about a unit vector *u* by an angle *θ*, for small enough *θ*, as a linear approximation according to

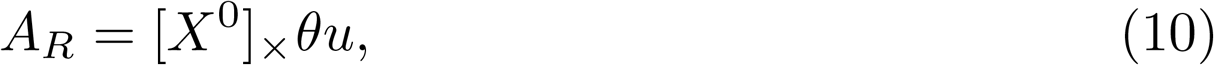

where *X*^0^ is the initial model used for the NMA computations, and[ · ]_×_ denotes the cross-product matrix,

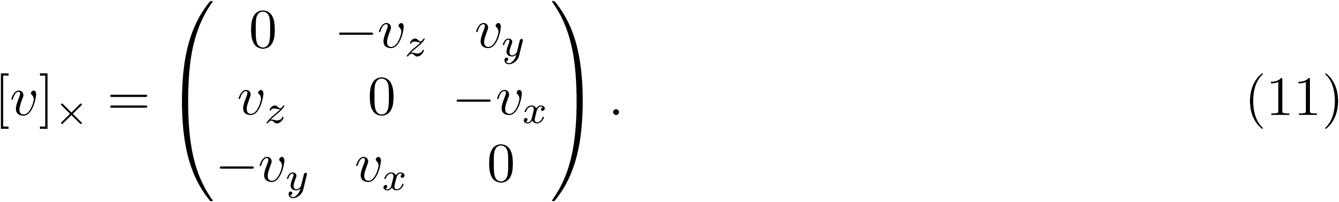

We determine the unknown normal mode amplitude Δ*α* at each iteration by minimizing the loss function *L* defined as

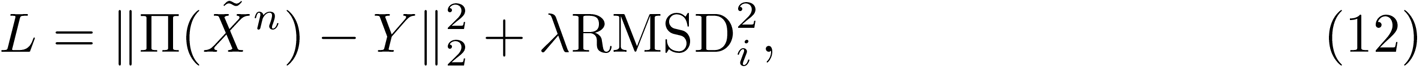

where *Y* is the experimental AFM image, Π is the smooth pseudo-AFM image function defined in Eq. 1, ‖ · ‖_2_ denotes the pixel-wise *l*_2_ norm, RMSD_*i*_ defines the *root-mean-square-deviation* between the initial model *X*^0^ and the fitted model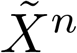, and *λ* is a parameter that controls the amplitude of the flexible deformation. The leftmost term in the loss function describes the similarity between the fitted atomic model and the experimental image. The right-most term is a regularization, which keeps the deformation amplitude of the initial model finite. Parameter *λ* controls the balance between data fitting and deformation amplitude and needs to be tuned carefully. Small *λ* values would attenuate constraints on the deformation amplitude and might result in model overfitting. At large *λ* values, the fitting would minimize the deformation amplitude and not deform the <u>m</u>odel sufficie<u>n</u>tly to achieve a good fit. We recommend scanning *λ* values between 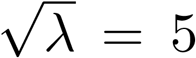 and 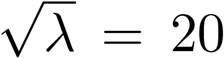 over multiple ima<u>ge</u>s (e.g., on a small subset of images) to find an optimal value. We used values of 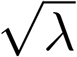 of 5, 20 and 10 for the experiments with EF2, FVa and TRPV3, respectively. To evaluate if the models are overfitted, one can use software such as Molprobity [41] that assesses their structural quality.

To obtain an analytical solution for Δ*α*, we perform the following linear approximation,

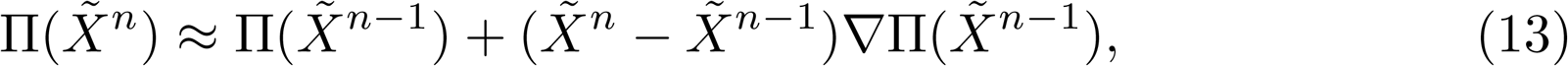

where ∇Π(*X*) is the gradient of the function Π(*X*) such as :

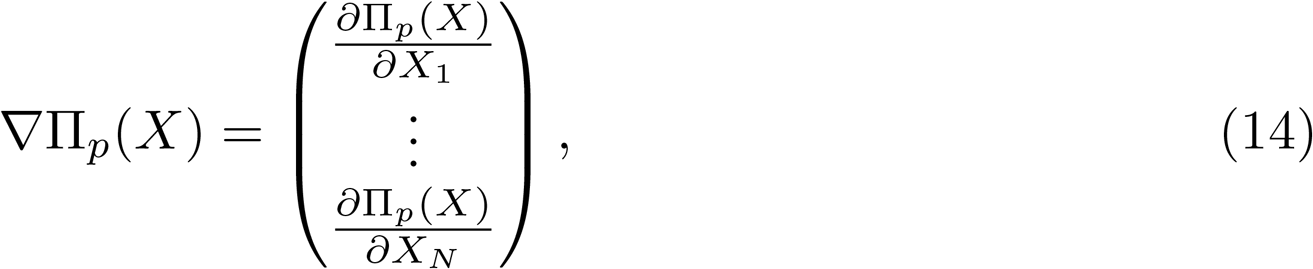

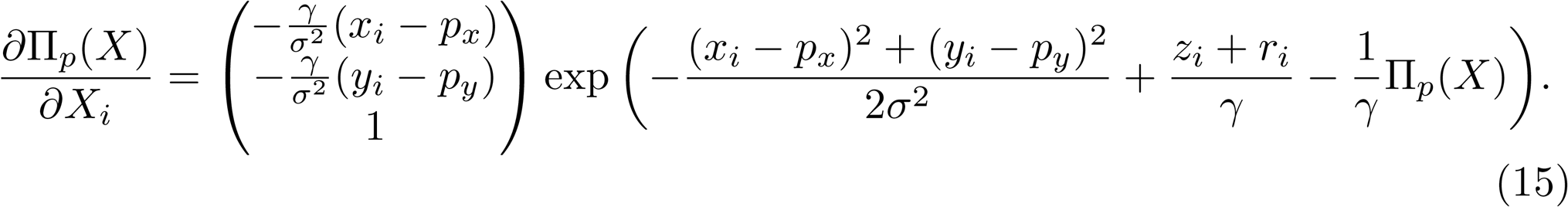

We also consider that the nonlinear extrapolation between each step is close to the linear transition :

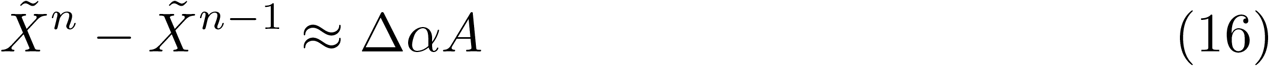

By using Eq. 16 and substituting Eqs. 9 and 13 in our loss function in Eq. 12, we obtain :

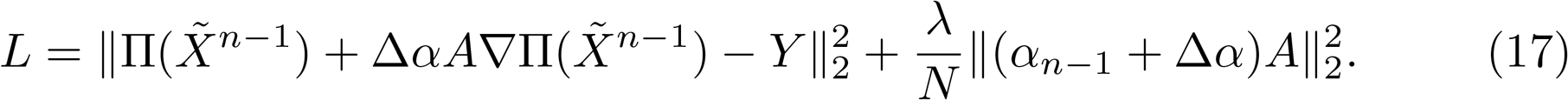

Finally, by equating partial derivatives of the Loss function with respect to the unknown amplitudes Δ*α* to zero,

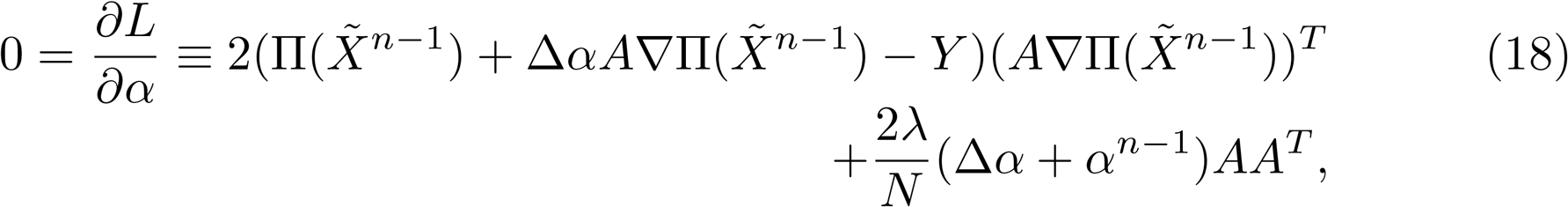

we obtain the final result:

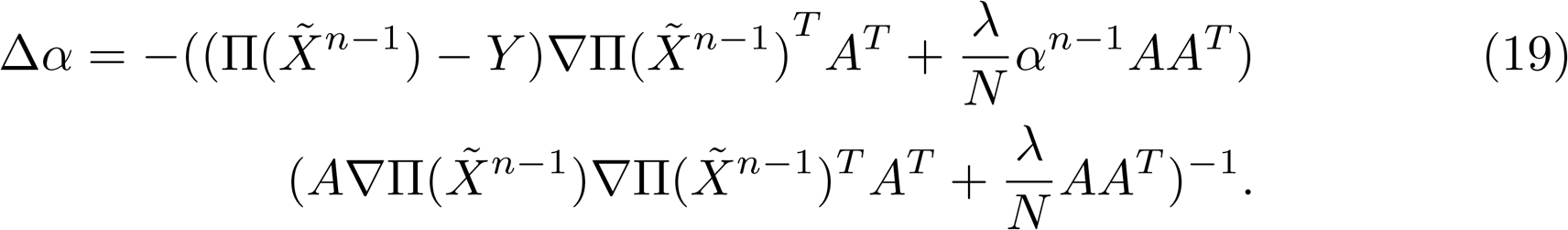

## Data availability

The source code of the method is freely available at https://gricad-gitlab.univ-grenoble-alpes.fr/GruLab/afmfit under the GPL-3.0 license. AFMfit comes as a Python package that can be installed with the *pip* package manager. We ship it with the NOLB normal modes package, which runs on Linux and MacOS operating systems.

The package provides code examples with GUI in the form of Jupyter notebooks. We also provide the reconstructed models for experiments with EF2, FVA and TRPV3, along with the synthetic AFM images of EF2 and the experimental AFM images of FVA. The raw HS-AFM images from SI movies 7 and 11 from [34] were kindly provided by Simon Scheuring. The original raw and corrected AFM images of FVA are deposited by Jean-Luc Pellequer at http://doi.org/10.5281/zenodo.11234725.

## Acknowledgements

We are very grateful to Simon Scheuring from Cornell University for providing us raw HS-AFM data of TRPV3. This work is supported by the French National Research Agency in the framework of the “Investissements d’avenir” program (ANR-15-IDEX-02).

**Supplementary information**

**Supplementary Video 1:** Reconstructed motion of FVA along the first principal component.

**Supplementary Video 2:** Reconstructed motion of FVA along the second principal component.

**Supplementary Video 3:** Fitting results of the first HS-AFM movie of TRPV3. Top left: the experimental HS-AFM movie of TRPV3. Bottom left: the pseudo-HS-AFM movie reconstructed from the fitted models. Top right: the difference between the experimental and the reconstructed HS-AFM movies. Bottom right: the fitted models.

**Supplementary Video 4:** Reconstructed motion of the TRPV3 tetramer along the second principal component for the first HS-AFM movie of TRPV3.

**Supplementary Video 5:** Reconstructed motion of the TRPV3 pentamer along the fifth principal component for the first HS-AFM movie of TRPV3.

**Supplementary Video 6** Fitting results of the second HS-AFM movie of TRPV3. Top left: the experimental HS-AFM movie of TRPV3. Bottom left: the pseudo-HS-AFM movie reconstructed from the fitted models. Top right: the difference between the experimental and the reconstructed HS-AFM movies. Bottom right: the fitted models.

